# Engineered muscle tissues with enhanced maturation enable the identification of clinically relevant rAAV capsids

**DOI:** 10.64898/2026.01.13.699271

**Authors:** Clémence Lièvre, Bérangère Robert, Giada Mainieri, Nicolas Jaulin, Marine Cotinat, Christian Mandrycky, David L. Mack, Oumeya Adjali, Caroline Le Guiner, Bodvaël Fraysse, Jean-Baptiste Dupont

**Affiliations:** Nantes Université, CHU Nantes, INSERM, TARGET, F-44000 Nantes, France; current affiliation: Department of Chemistry, Materials and Chemical Engineering, Politecnico di Milano, Milan, Italy; Institute for Stem Cell and Regenerative Medicine, Department of Rehabilitation Medicine, University of Washington, Seattle, WA, USA; Department of Bioengineering, University of Washington, Seattle, WA, USA

**Keywords:** Engineered muscle tissues, iPSCs, in vitro models, gene therapy, AAV vectors, tissue engineering

## Abstract

Developing in vitro models that recapitulate both the structure and function of native human tissues is crucial for a better understanding of pathophysiology and for improving the reliability of preclinical studies. Here, we demonstrate that engineered muscle tissues derived from human pluripotent stem cells can serve as an in vitro platform for gene therapy. Recombinant vectors derived from the adeno-associated virus transduce engineered muscle tissues with high efficiency and in a dose-dependent manner, allowing long term assessment of transgene expression in a human cellular context. We next used this model to conduct a comparative analysis of 8 natural AAV capsids and showed that their relative efficiency depends on engineered muscle tissue maturation level. In more mature tissues subjected to uniaxial mechanical stretch, AAV9 performed better, which is reminiscent of its high clinical potential in patients with neuromuscular disorders. Finally, our model also confirmed the higher efficiency of artificial MyoAAV variants specifically developed to have an improved muscle transduction. Altogether, this work demonstrates the potential of human engineered muscle tissues in the preclinical testing of AAV vectors, paving the way for the development of personalized gene therapy platforms.

## INTRODUCTION

Preclinical development of gene therapy strategies for neuromuscular disorders (NMDs) integrates vector design, production in cellular systems and the comprehensive evaluation of efficacy and safety. Thus far, the latter has employed animal models such as rodents, dogs and non-human primates, as their skeletal muscles share strong physiological similarities with humans. In animal models of Duchenne muscular dystrophy (DMD), Limb-girdle muscular dystrophy or X-linked myotubular myopathy (XLMTM), systemic injection of recombinant vectors derived from the adeno-associated virus (rAAV) expressing therapeutic transgenes has led to tremendous phenotypic improvements (^1–4^). Complete and durable disease rescue has become a common outcome in preclinical muscle gene therapy, which led to the initiation of clinical trials in the years 2018 – 2020. More than 5 years later, and in spite of promising intermediary results observed after injection, the trend is rather toward a lower overall efficacy in patients than that achieved in animal models (^5^). More importantly, while systemic rAAV injection was safe and well-tolerated in animal models of NMDs, several patients developed serious adverse events including thrombotic microangiopathy due to complement activation (^6^), renal impairment, myocardial injury and myositis (^7,8^). These toxic effects have been linked with acute immune reactions against the AAV capsid or the transgene and have led to clinical trial discontinuation. Anti-AAV immunity is known to be dose-dependent (^9^) and is likely exacerbated by the high vector doses required to reach therapeutic efficacy in patients with NMDs. Altogether, these observations underscore the need for next-generation AAV vectors with an improved therapeutic index, but also for innovative experimental models that better recapitulate patient physiology and are compatible with the high throughput screening experiments required to identify the optimal gene therapy product and dose regimens.

This study aims to develop an *in vitro* preclinical testing platform for muscle gene therapy by harnessing the potential of human pluripotent stem cells and tissue engineering. Disease modelling has undergone a revolution since the discovery of induced pluripotent stem cells (iPSCs) carrying the genetic background of the patients (^10^). Thanks to directed differentiation protocols giving rise to skeletal muscle myotubes in a dish, research is progressively moving toward modelling of NMD phenotypes using the patient’s own cells. Differentiation of iPSCs from patients with DMD leads to impaired myotube formation, dysregulation of gene expression, release of creatine kinase (CK) into the media as a sign of cellular damage, Ca^2+^ hyperexcitability and contraction defects (^11–13^). In addition, iPSC-derived models are expandable and can be used in drug discovery studies to evaluate the efficacy of pharmacological compounds (^13,14^), discover new therapeutic candidates (^15^), or validate gene transfer/editing strategies (^11,12^). An important limitation of these new models is linked to the low cellular maturation obtained after differentiation in standard 2D monolayer cultures, at both structural and functional levels. Myotubes derived from iPSCs are thin and spindly, show heterogeneous striation and multinucleation patterns together with uncontrolled contractions. We and others have developed engineered muscle tissues (EMTs) grown between two posts in defined 3D hydrogel environments to model the molecular and cellular phenotypes of NMDs such as Duchenne and limb-girdle muscular dystrophies (DMD and LGMD, respectively) (^16–18^) and laminopathies (^19,20^). Switching to such 3D “tissue-like” configuration allows to mimic the longitudinal organization of skeletal muscles, but iPSC-derived myotubes never progress to *bona fide* muscle fibers. Three-dimensional muscle constructs derived from primary or immortalized myoblasts are known to be highly sensitive to the biophysical cues naturally present in the niche during development. Mechanical conditioning of engineered constructs containing C2C12 murine muscle cells leads to increased expression of Myosin heavy chain proteins (^21^). Human bioartificial muscles submitted to stretch-relaxation cycles showed increased myofiber diameter and area (^22^). Thus, we posit that mature EMTs can provide a relevant model for preclinical gene therapy and serve as a complement to animal experimentation.

In the present study, we demonstrated the feasibility of rAAV-mediated gene transfer in iPSC-derived EMTs, and we assessed the impact of tissue maturation on transduction efficiency. Using a reporter transgene, we determined the optimal transduction parameters, the transgene expression kinetics and the minimum effective dose. We next introduced the MyoStretch maturation platform, a custom-made 3D culture module designed to implement constant uniaxial mechanical stretch during EMT differentiation. We showed that mechanical stimulation improved EMT maturation, promoting better myotube alignment and lower dependence to extracellular calcium. Importantly, comparison of eight natural AAV serotypes showed that EMTs with a higher level of maturation were better at discriminating capsid 9, which is currently injected in NMD patients enrolled in clinical trials in relation to its good muscle tropism. Additionally, our EMT platform successfully identified the most efficient capsid among recently developed myotropic variants, further supporting the relevance of iPSC-derived models for rAAV vector preclinical development.

## RESULTS

### Addition of rAAV vectors in iPSC-derived EMTs leads to efficient transgene transduction

In order to evaluate the efficiency of rAAV vectors transduction in EMTs, we first used iPSCs derived from a healthy donor. iPSCs were subjected to directed differentiation into the skeletal muscle lineage using a transgene-free and serum-free protocol (^23^). Myogenic progenitors obtained after 28 to 35 days of differentiation were included in a gel composed of Matrigel® and fibrin to form EMTs between two pillars, as previously published (^16,19^). A reporter AAV vector of serotype 6 was produced to express the murine secreted alkaline phosphatase (mSEAP) downstream of the muscle-specific Spc5-12 promoter (Figure 1A) (^24^). The mSEAP transgene was designed to allow mSEAP protein release from transduced cells, that can be used for longitudinal assessment of transduction efficiency. Using a specific design of experiments (DOE) approach that enables to assess the influence of multiple experimental factors in a minimal number of combinations, we were able to test the influence of 3 factors simultaneously: the mode of vector delivery (gel inclusion *vs.* addition in medium), the dose (1E+4vg/cell *vs.* 1E+5 vg/cell) and the culture media maintained during the experiment (proliferation media *vs.* differentiation media) (Figure S1A). Transduction efficiency was assessed by measuring mSEAP expression in the culture supernatant at different days post-transduction by luminometry and by measuring mSEAP expression at day 10 post-transduction by RTqPCR. Briefly, we showed that inclusion of the rAAV6 reporter vector directly in the hydrogel led to a higher transduction efficiency than addition into the media. Similarly, transduction was more efficient when EMTs were placed in differentiation media 3 days after transduction, compared to EMTs maintained in proliferation media (Figure S1B). On the other hand, the dose of vector seems to only have low impact on transduction efficiency (Figure S1C). Therefore, for the next steps of the study, rAAVs were added directly into the hydrogel during EMTs formation and differentiation protocol into myotubes was launched after transduction.

**Figure 1:**
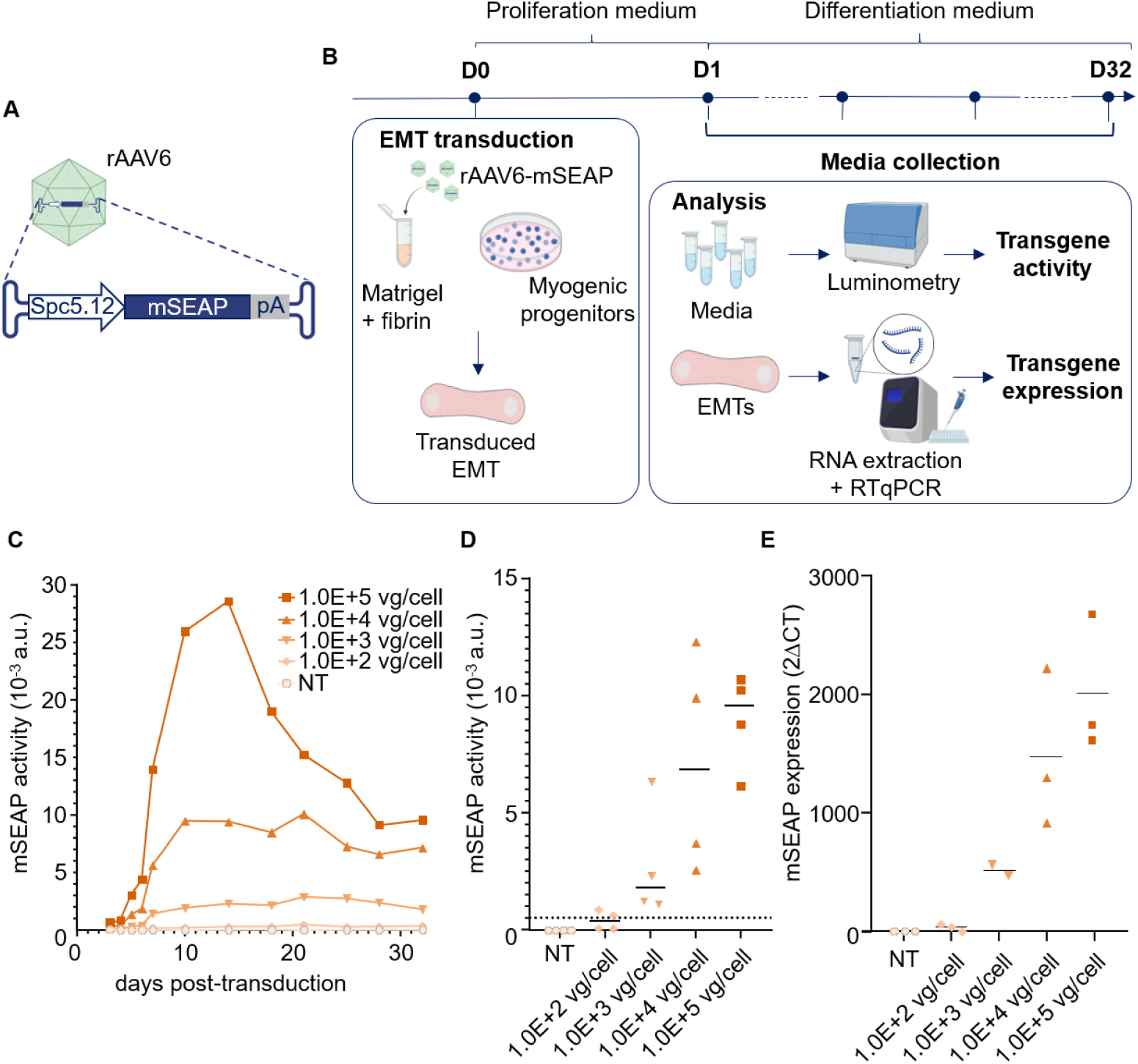
Inclusion of AAV vectors in engineered muscle tissues leads to high transduction efficiency. (A) Reporter rAAV vector including the Spc5.12 promoter, mSEAP reporter cassette in a serotype 6 capsid (created using Biorender). (B) Experimental design to assess transduction efficiency. The rAAV6-Spc5.12-mSEAP reporter vector was included directly into the hydrogel during EMT formation and supernatants were collected every 2 days at each media change. At the end of experiment, mSEAP transgene activity was measured by luminometry on media samples and transgene expression was assessed by RTqPCR on EMT RNA (created using Biorender). (C) Kinetics of mSEAP activity after EMT transduction with decreasing doses of vector. NT: non-transfected control. Each point represents the mean of 4 EMTs. (D) mSEAP activity at day 32, with a threshold at 500% above the negative control (NT). Each point represents an independent EMT (n=4 per group). Bars indicate the median value. (E) Relative quantity of mSEAP after RNA extraction from EMTs at day 32 post-transduction and normalization with *UBC* expression. Each point represents an independent EMT (n=2 or 3 per group). Bars indicate the median.

Then, we aimed to determine the minimum effective dose of rAAV vector required to detect transgene expression in our experimental settings. The reporter rAAV6-Spc5.12-mSEAP vector was added in EMTs derived from iPSCs during hydrogel formation, at decreasing doses ranging from 1E+5 vg/cell to 1E+2 vg/cell. Aliquots of culture media were collected every other day and transgene expression was analyzed by luminometry at the end of the experiment (Figure 1B). Transgene expression was detected as soon as the 1.0E+3 vg/cell dose and as early as Day 4 post transduction. It then increased until 10 days post-transduction before reaching a plateau which was stable until more than 30 days post transduction (the longest time point evaluated). The expression profile was comparable with the 1.0E+4 vg/cell dose, but was quite different with the highest 1.0E+5 vg/cell dose: with this high dose, mSEAP activity peaked at D10 and then steadily decreased until D28 post transduction, suggesting vector-mediated toxicity or EMT saturation with short-lived rAAV genome copies (Figure 1C). To compare transduction at the experimental endpoint, a threshold was arbitrarily set at 500% of the background mSEAP activity obtained at D32 with non-transduced EMTs. The minimum dose allowing consistent detection of mSEAP activity above the threshold was clearly 1.0E+3 vg/cell, which was quantified as 100-fold higher than the non-transduced negative control (Figure 1D). This result was confirmed at the transgene mRNA level by RTqPCR at D32 post-transduction (Figure 1E). In the next steps of our study, rAAV vectors were thus used at a dose of 1E+3 vg/cell to better see any differences when comparing, and experiments were carried out for 10 days, where transgene expression reaches a plateau.

### Capsid comparison in iPSC-derived EMTs highlights limited maturation level

Using the experimental conditions optimized above, we next compared the transduction efficiency of 8 natural rAAV capsids (rAAV1,2,3b,5,6,8,9 and 10) at 1E+3vg/cell in standard EMTs generated from healthy iPSCs and cultured between two posts (Figure 2A-B). mSEAP transgene expression and enzymatic activity were monitored overtime. The overall kinetics were similar with all capsids and consistent in a regular increase up to 10 days post-transduction (Figure 2C). Transgene expression at Day 10 was assessed by RTqPCR after RNA extraction from transduced EMTs. In this experimental context, rAAV6,2 and 1 allowed the highest transgene expression levels. On the other hand, rAAV serotypes 8 and 9, known to be the most effective serotypes *in vivo* in muscle cells (^25^) and currently used in several clinical trials (^4^), were 10 to 100 times less effective than rAAV6. We compared transduction efficiency between each serotype and significant differences (Kruskal-Wallis test, *p*-value <0.05) were observed between rAAV6 and rAAV5 (*p*=0.0048), rAAV6 and rAAV9 (*p*=0.0169), rAAV2 and rAAV5 (*p*=0.0119), and rAAV2 and rAAV9 (*p*=0.0444) (Figure 2D). Although rAAV6 and rAAV2 have been used in early *in vivo* studies in DMD animal models (^26,27^), these capsids are also known to be among the most efficient for *in vitro* transduction experiments. In myogenic progenitors differentiated in 2D culture conditions, we confirmed that rAAV6 and rAAV2 led to significantly higher mSEAP activity compared to all other serotypes (one-way ANOVA, *p*-value<0.05) (Figure S2). Altogether, these results indicate that the iPSC-derived EMT platform still mostly reproduces *in vitro* 2D culture conditions rather than a true skeletal muscle tissue state, underscoring the need to improve maturation strategies.

**Figure 2:**
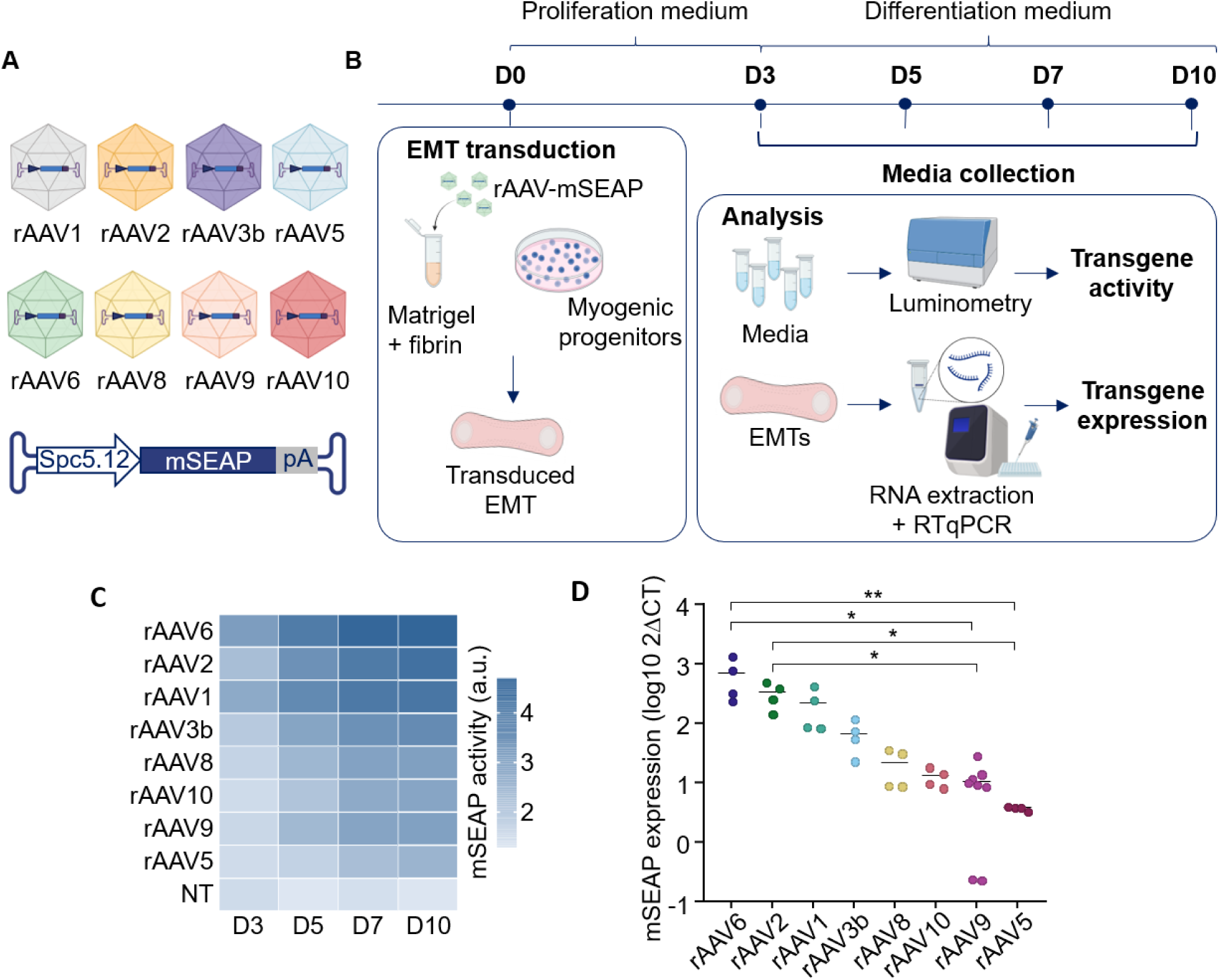
Capsid comparison in iPSC-derived EMTs highlight limited maturation level. (A) Reporter Spc5.12-mSEAP expression cassette in 8 natural AAV capsids (created using Biorender). (B) Experimental design to assess transduction efficiency. The reporter vectors were added into the hydrogel during EMT formation and supernatants were collected at D3, D5, D7 and D10. At the end of experiment, mSEAP activity was measured by luminometry on media samples and transgene expression by RTqPCR on EMT RNA (created using Biorender). (C) Kinetics of mSEAP activity increase after transduction of EMTs with different serotypes of rAAV-Spc5.12-mSEAP. A darker blue box represents a higher median luminescence value (N = 5 to 6 EMTs per serotype), represented in arbitrary units (a.u.) with logarithmic scale. (D) Relative mSEAP expression after RNA extraction from EMTs at 10 days after transduction with 8 different natural rAAV serotypes and normalization with *UBC* expression. Each point represents the value obtained with one EMT (n=4 or 8 per group). Bars indicate the median. Statistics: Kruskal-Wallis test (*p-value<0.05, **p-value<0.01).

### Increased cellular density and mechanical stretch improve EMT maturation

We next sought to optimize EMT maturation to better recapitulate the structure and function of native skeletal muscle tissues. For this purpose, we designed MyoStretch, a home-made device able to implement constant uniaxial mechanical tension allowing to stretch EMT during differentiation (Figure 3A). In this configuration, the EMT hydrogel is formed between a rigid pillar and a flexible pillar, which can be moved apart from each other using a screw. In preliminary experiments, we compared the contraction of EMTs stretched at +5, +10 and +15% of their initial length with unstretched EMTs. We observed that 10 and 15% stretch regimens led to significant increases in contraction amplitude (Figure S3). For the rest of the study, we selected the 15% stretch condition, which provides the highest amplitude while allowing for additional elongation later in the protocol. In addition, we increased the number of cells per EMT with 2.0E+6 myogenic progenitors, in an attempt to increase the density of contractile cells in the hydrogel.

**Figure 3:**
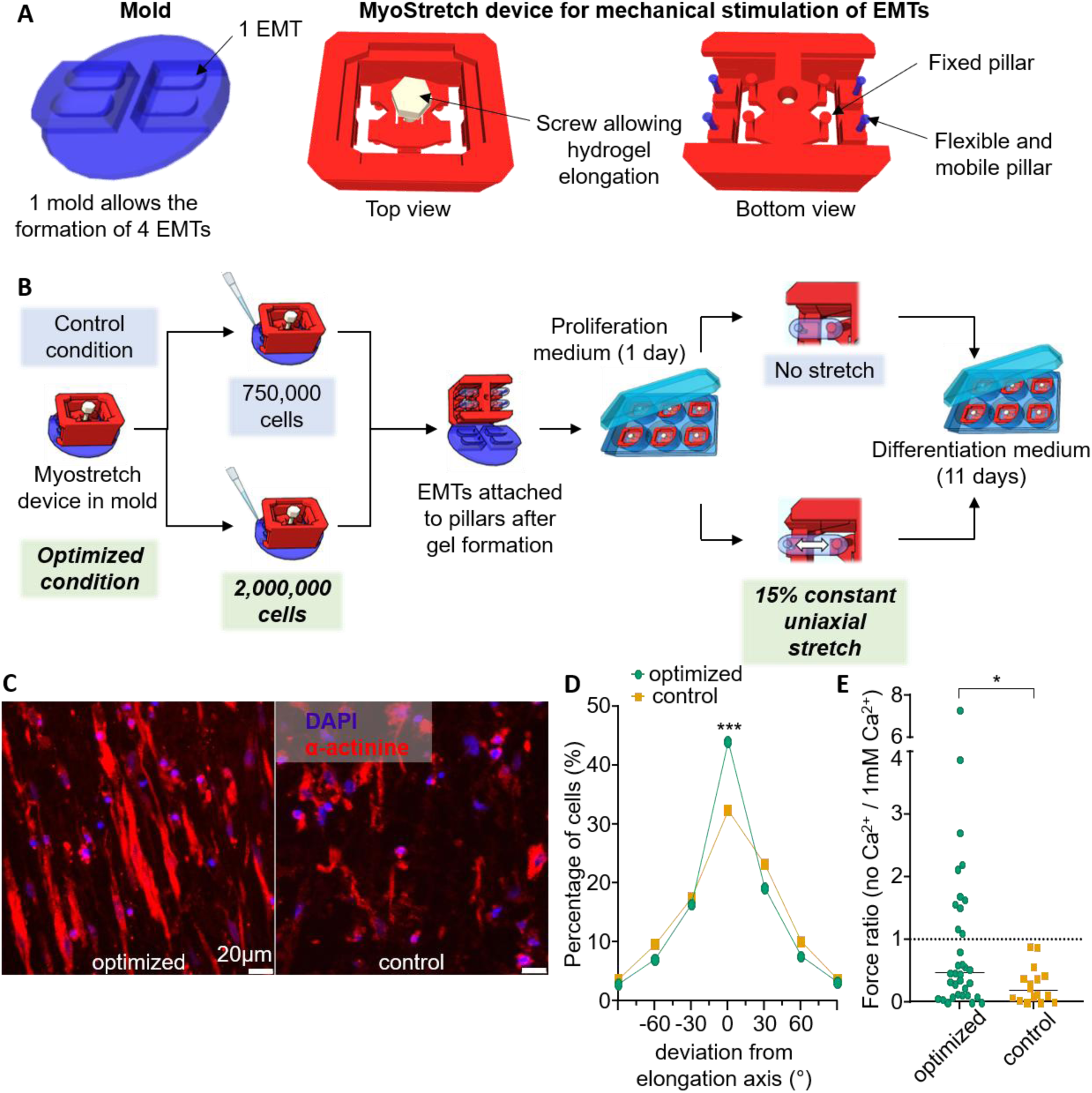
Increased cellular density and mechanical stretch improve EMT maturation. (A) Design of a custom-made Myostretch device for mechanical elongation of EMTs. (B) Experimental design of EMT formation and differentiation in control and optimized maturation conditions. (C) Confocal microscopy representative pictures of EMT cryosections subjected to immunofluorescence against α-actinin (red). Nuclei were stained with DAPI (blue). Scale bars: 20µm. (D) Quantification of myotube alignment from immunofluorescence pictures from optimized and control EMTs. Each point indicates the mean of 86 values for the optimized condition and 37 values for the control. Five images were taken and analyzed per EMT. (E) Ratio of contractile forces generated by EMTs in the optimized *vs.* control conditions in the absence of Ca^2+^ or with 1mM Ca^2+^. Each point represents the values obtained for one EMT (n=34 for the optimized condition, n=16 for the control). Bars indicate the median of each group. Statistics: Mann-Whitney U-test (*p-value<0.05).

Ultimately, we compared the morphology and function of EMTs generated in optimized conditions (*i.e.* 2.0E+6 cells per tissue and 15% stretch) with EMTs produced with a standard protocol (0.75E+6 cells per tissue, 0% stretch) (Figure 3B). We first assessed cell alignment within EMTs by fluorescence microscopy after staining of the sarcomeric protein α-actinin. In optimized tissues, α-actinin-positive myotubes were visibly larger and oriented along the elongation axis (Figure 3C). We quantified the percentage of cells oriented at different angles and observed that the optimized maturation condition led to a significant increase in the proportion of aligned myotubes (+/- 15° aligned with the elongation axis: 44% in the optimized condition *vs.* 30% in the control, Mann-Whitney U test: p = 0.0004) (Figure 3D). At the functional level, we observed no improvement of the contraction force amplitude in optimized compared to non-optimized EMTs (data not shown). Thus, we evaluated EMT contraction dependence on extracellular calcium, as mature fibers mobilize intracellular stores while immature tissues rely on extracellular calcium influx (^28^). The ratio between the force obtained without the addition of extracellular calcium and the force obtained when EMTs were incubated with 1mM of calcium in the culture medium was significantly higher in the “optimized” condition than in the “control” one (optimized: median ratio = 0.47, control: median ratio = 0.19, Mann-Whitney U test: p = 0.03) (Figure 3E). This indicates that overall, our optimized maturation protocol leads to EMTs that are less dependent on extracellular calcium and more capable of contracting by mobilizing their internal stock of calcium.

### Capsid comparison outcome depends on EMT maturation

Our working hypothesis is that EMTs with an improved level of maturation will better predict gene therapy efficiency in preclinical and then clinical studies. To assess this point, the transduction efficiency of the 8 natural rAAV capsids previously tested in non-optimized EMTs was re-evaluated in EMTs with an optimized level of maturation. Similar to previously, reporter rAAV vectors were added directly in the hydrogel at 1E+3 vg/cell during the formation of EMTs. Mechanical stretch at 15% was added on the following day, upon induction of differentiation. Longitudinal monitoring of transgene activity in optimized maturation conditions resulted in a comparable profile to that obtained in non-optimized tissues, with a steady increase for 10 days (Figure 4A). Nevertheless, the 8 natural capsids performed differently when compared to one another, and rAAV9 emerged as the most efficient serotype in more mature EMTs, which aligns well with its clinical use in patients with NMDs. The expression of the mSEAP transgene was measured at day 10 post-transduction by RT-qPCR and confirmed that rAAV9 resulted in the highest transgene expression in optimized EMTs, even if the difference was non-significant (Kruskal-Wallis statistics, p-value>0.05) (Figure 4B). Altogether, these results show that EMTs with an optimized level of maturation better recapitulate the known relative efficiency of rAAV capsids observed in skeletal muscles of NMD animal models. These observations then prove that capsid comparison outcome depends on EMT maturation level.

**Figure 4:**
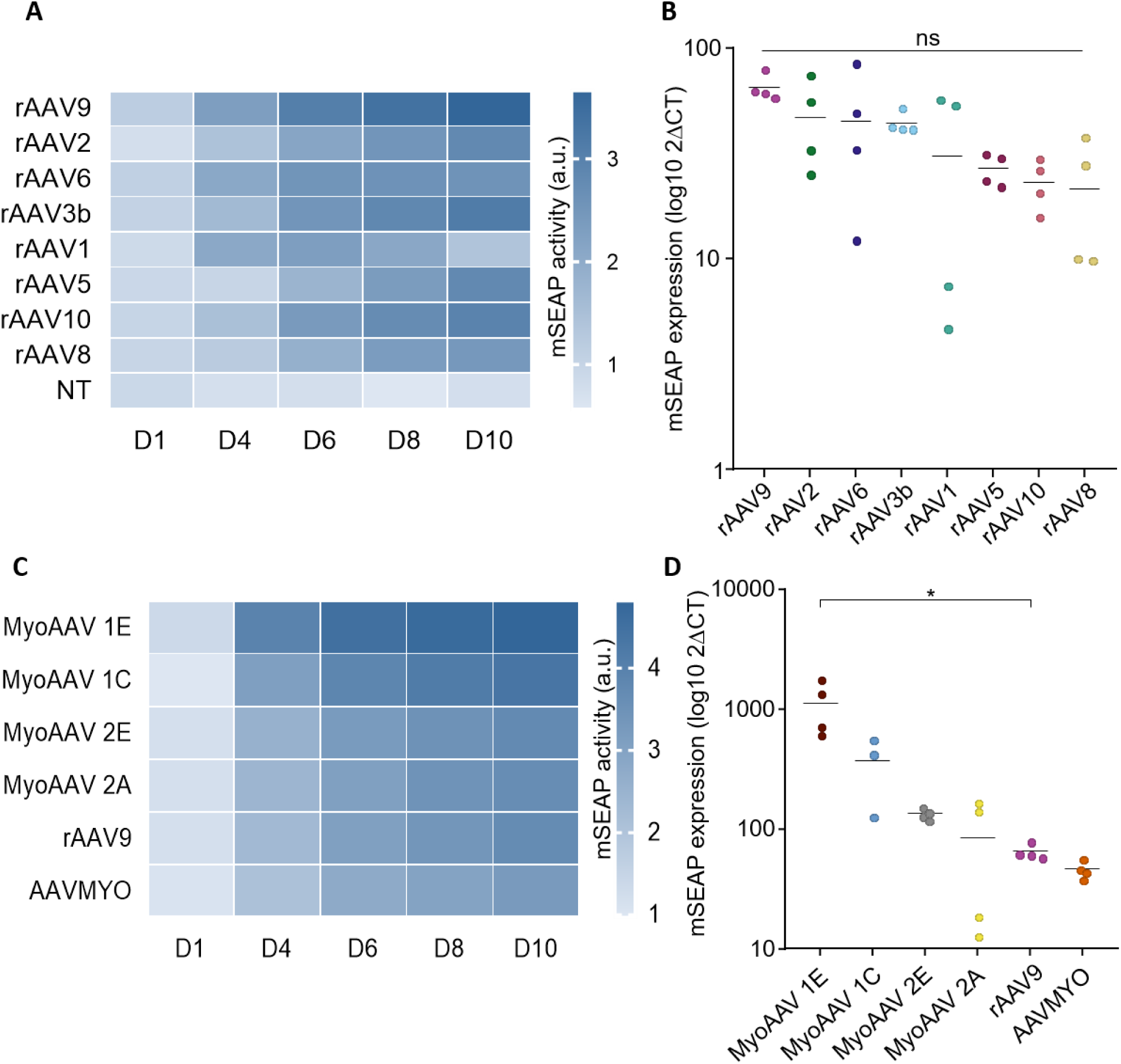
Capsid comparison outcome depends on EMT maturation. (A) Kinetics of mSEAP activity after transduction of optimized EMTs with different serotypes of rAAV-Spc5.12-mSEAP. A darker blue box represents a higher median luminescence value, represented in arbitrary units (a.u.) with logarithmic scale (N = 2 to 4 EMTs per serotype). (B) Relative expression of mSEAP after RNA extraction from EMTs at 10 days after transduction with different rAAV serotypes, and normalization with *UBC* expression. Each point represents an independent EMT (n=4 per group). Bars indicate the median value. (C) Kinetics of mSEAP activity after transduction of optimized EMTs with natural and artificial muscle-tropic capsids of rAAV-Spc5.12-mSEAP. The median luminescence values are represented in arbitrary units (a.u.) with logarithmic scale (N = 2 to 4 EMTs per serotype). (D) Relative expression of mSEAP after RNA extraction from EMTs at 10 days after transduction with different muscle-tropic rAAV serotypes, and normalization with *UBC* expression. Each point represents an independent EMT (n=4 per group). Bars indicate the median. Statistics: Kruskal-Wallis test (n.s.: not significant; * p-value < 0.05).

Ultimately, we assessed the ability of our iPSC-derived EMT platform to predict the higher efficiency of previously published rAAV variants designed to better target skeletal muscle tissues. The Spc5-12 mSEAP reporter cassette was packaged in five different myotropic capsids derived from the rAAV9 serotype and previously published by two different groups: the original AAVMYO capsid (^29^), and four MyoAAV capsids (MyoAAV 1C, MyoAAV 1E, MyoAAV 2A and MyoAAV 2E) (^30^). The efficiency of myotropic vectors was compared to that of rAAV9 in mature EMTs, by monitoring mSEAP activity by luminometry for 10 days post-transduction (Figure 4C). Interestingly, we observed that all MyoAAV variants led to comparable or higher mSEAP activity at Day 10 post-transduction. MyoAAV 1E was the most efficient and resulted in a 14-time higher activity than AAV9. Transgene expression was analyzed by RTqPCR, which confirmed the higher efficiency of all MyoAAV variants. The efficiency of the AAVMYO variant, which was primarily designed to de-target the liver, appears not significantly different – and even slightly lower – than that of AAV9 (Figure 4D).

## DISCUSSION

In this study, we demonstrated that bioengineered muscle tissues derived from human iPSCs can be used as a preclinical platform to evaluate the *in vivo* efficacy of gene therapy vectors. In term, they could provide a complement to animal experimentation at specific steps of the rAAV drug discovery pipeline. In the context of NMDs, animal models have been instrumental to decipher the disease mechanisms at a molecular level and to prove the feasibility of global preclinical strategies such as gene therapy. Over the years, they have however showed several limitations inherent to interspecies differences. For example, in the *mdx* mouse model of DMD, compensatory mechanisms have been highlighted, such as upregulation of the utrophin gene (^31^), presence of long telomeres (^32^) or expression of cytidine monophosphate sialic acid hydroxylase (*Cmah*) (^33^). These murine-specific mechanisms likely contribute to the milder phenotype of *mdx* mice. Furthermore, the GRMD dog model exhibits the symptoms of the disease in a heterogeneous manner and even displays phenotypes rarely observed in patients, such as dysphagia (^34^), thereby limiting the translational value of discoveries and therapeutic testing performed in these models for humans.

Assuming that models in a human genetic background would be more specific to recapitulate the mechanisms of NMDs, we developed a human-based preclinical testing platform for rAAV muscle gene therapy. We used our expertise in myogenic differentiation of human iPSCs to generate the large batches of myogenic progenitors required for the formation of EMTs. The transgene-free differentiation protocol used in the present study is known to generate alternative cell types (^35^), hence requiring further purification. We sorted myogenic progenitors expressing the surface marker ERBB3, which was specifically identified in mononucleated cells from fetal muscles (^36^). As previously shown by other groups, cell sorting substantially improved the quality of myotube differentiation (data not shown) and allowed to eliminate a large fraction of negative cells that could have reduced EMT formation efficacy. In our study, we also used a custom-made culture module to add mechanical stretch during EMT differentiation, in an attempt to reproduce the mechanical tension supported by skeletal muscle cells during development (^37^). Mechanical conditioning has a long track record in improving the maturation of skeletal muscle cells in 2D cultures (^38,39^) but also in the first generations of artificial muscle based on primary cells (^21,22^). At molecular and cellular levels, the application of 5% to 20% stretch regimens led to increased synthesis of contractile proteins, myotube hypertrophy and alignment and activation of anabolic signaling pathways (^40^). Using MyoStretch, our study further supports beneficial effects of mechanical stretch on the maturation of EMTs derived from iPSCs, both on myotube alignment and functional dependence to calcium. Further characterization will be required to precisely determine the impact of constant uniaxial stretch on gene expression, remodeling of myotube architecture and their consequences on the contractile function. Additional cues could be implemented and help further improve the maturation level of iPSC-derived EMTs, such as electrical stimulation to mimic neural innervation or addition of other cell types (*i.e.* fibroblasts and endothelial cells). Nevertheless, MyoStretch allows the development of more mature EMTs, supporting their use as a relevant preclinical model.

In non-optimized conditions, our results indicate that EMTs have the potential to serve as a preclinical testing platform for rAAV gene therapy. Transgene activity could be monitored for more than 1 month and its expression was still detectable at 32 days post transduction (the longest time-point evaluated), which to our knowledge has never been previously achieved in an *in vitro* human cellular system. On this time window, transgene activity also appears stable and dose-dependent. With the highest M.O.I. (1.0E5 vg/cell), it progressively declined after a high peak of activity, suggesting cellular toxicity or saturation of endogenous genome processing and/or gene expression machineries, leading to the silencing or elimination of specific genome forms (^41^).

Attempts at improving the therapeutic index of rAAV vectors are currently being made through the guided development of more efficient genomic elements or new capsids targeting muscle cells with higher specificity and minimal transduction in off-target organs (^29,30^). This results in a high number of potential vector candidates which needs to be tested and compared prior to possible clinical translation. Capsid DNA shuffling or barcode ligation allow to multiplex these candidate vectors and quantify their transduction efficiency by high-throughput sequencing in a minimal number of *in vivo* experiments (^30,42^), but this strategy remains difficult to implement. We propose that EMTs derived from iPSCs could serve as a relevant *in vitro* testing platform. We successfully compared 8 natural and 5 artificial capsids of rAAV packaging an mSEAP expression cassette. Interestingly, we showed that transduction efficiency was impacted by the level of EMT maturation, and that more mature EMTs were better at predicting the efficiency of capsids with known efficiency in skeletal muscles *in vivo*, such as AAV9 and several myotropic variants. The impact of mechanical stretch on EMT permissiveness to rAAV remains to be precisely characterized and may affect multiple steps of the cellular transduction process, from vector internalization, intracellular trafficking or epigenetic regulation of transgene expression. Most strikingly, the parallel comparison of 4 MyoAAV variants resulted in a similar ranking in iPSC-derived EMTs as the one obtained with primary human myotubes in the initial study (^30^). Our EMT model derived from iPSCs remains significantly more expandable than primary muscle cells and is therefore able to evaluate substantially more vector candidates in parallel experiments. Thus, the consistency observed between the two model systems strongly suggests that high-throughput screening experiments conducted in EMTs will have a high preclinical relevance and predictive power.

Altogether, the present study demonstrates the preclinical potential of iPSC-derived EMTs for muscle gene therapy and highlights the importance of improving their level of maturation. Continuous efforts to bioengineer more mature muscle tissues is therefore indispensable for the successful deployment of this model in next generation rAAV drug discovery pipelines.

## MATERIALS AND METHODS

### Ethical statement

The Healthy UC3-4 line was generated from a human adult male donor with no history of skeletal muscle disease. Urine sample was used to isolate urine stem cells (UC) that were expanded and reprogrammed to generate iPSCs at the University of Washington, Seattle (USA) (^16,43^). The donor gave informed consent, as required by the Institutional Review Board (IRB). The cell line was extensively characterized, expanded and shared with the TaRGeT INSERM laboratory at Nantes Université through a dedicated Material Transfer Agreement. The experiments presented in this study fall into the Category 1A described in the Guidelines for Stem Cell Research and Clinical Translation of the International Society for Stem Cell Research (ISSCR, https://www.isscr.org/guidelines).

### iPSC maintenance

All cell culture experiments were performed at 37°C and 5 % CO2 in a standard tissue culture incubator. Master Cell Banks (MCB) were generated at passage 34. Cells were thawed at 37°C and plated as clusters on Matrigel (1:60 dilution, Matrigel® matrix, Corning, #354234) after spinning 3 min at 300 g. Amplification was made in mTeSR Plus culture medium (mTeSR™ Plus, Stemcell technologies, #100-0276) supplemented in 10 μM ROCK inhibitor (Y-27632 ROCK inhibitor, Selleckchem, #S1049) for the first day. Fresh media without ROCK inhibitor was renewed every other day. Abnormally looking cell clusters were manually removed every other day, until cells reached 70 to 80 % confluency. Cells were then passaged with Versene (Versene solution, Thermo Fisher, #15040066). Working Cell Banks (WCB) were generated from the MCB after two serial passages. Cells were detached with Versene and cell scraper, spun down 3 min at 300 g, and clusters were gently resuspended in Cryostor (Cryostor CS10 cryopreservation medium, Stemcell technologies, #07959) freezing media for long term cryopreservation in liquid nitrogen.

### iPSC myogenic differentiation

Directed differentiation of iPSCs was carried out with a succession of five defined media, as published previously (^23^). Single cells were seeded on Matrigel-coated 6-well plates (1:60 dilution) as at a density of 25,000 cells / cm², and were amplified for 2 – 3 days in mTeSR Plus media before differentiation induction. ROCK inhibitor was added upon seeding but removed by a fresh media change after one day. The cells were then incubated in five successive differentiation media based on Dulbecco’s Modified Eagle’s Medium (DMEM) / F12 (DMEM/F12, Thermo Fisher, #11320033) supplemented with Non-Essential Amino Acids (NEAA, Thermo Fisher, #11140035) and defined cytokines and growth factors, as detailed below:

- Media 1 (3 days): Insulin transferrin selenium (ITS, Thermo Fisher, #51500056) 1X + 3 μM CHIR99021 (Stemcell technologies, #72054) + 0.5 μM LDN (StemMACS™ LDN-193189, Miltenyi Biotec, #130-103-925)
- Media 2 (3 days): Insulin transferrin selenium (ITS) 1X + 3 μM CHIR99021 + 0.5 μM LDN + basic Fibroblast Growth Factor (bFGF) at 20 μg / ml (recombinant human FGF2, R&D systems, #233-FB-025)
- Media 3 (2 days): 15 % Knockout serum replacement (KSR, Thermo Fisher, #10828010) + 0.5 μM LDN + bFGF at 20 μg / ml + Hepatocyte Growth Factor (HGF, R&D systems, #294-HG-005) at 10 μg / ml + Insulin-like Growth Factor 1 (IGF, R&D systems, #291-G1-200) at 2 ng / ml.
- Media 4 (4 days): 15 % KSR + IGF-1 at 2 ng / ml
- Media 5 (16 days): 15 % KSR + IGF-1 at 2 ng / ml + HGF at 10 μg / ml

Cells were collected between day 28 and day 32 after incubation with a combination of Trypsin-EDTA (Sigma-Aldrich #T4174) and collagenase IV (Thermo Fisher, #1710419) at 50 U / μl for 10 min at 37 °C, mechanical dissociation and passage through a 70 μm cell strainer to remove debris and extracellular matrix. Myogenic progenitors thus obtained are then directly seeded in Matrigel-coated 15cm plates for amplification and sorting.

### Myogenic progenitor maintenance

Myogenic progenitor cells directly isolated from the myogenic differentiation of iPSCs were seeded on Matrigel-coated plates (1:60 dilution) at 5 000 cells/cm² after spinning 5 min at 300 g. A commercial skeletal muscle growth medium (SkGM2®, Lonza, CC-3245) was used for amplification. Fresh media was renewed every other day. When cells reached a 80 to 90 % confluence, they were sorted by Fluorescence Activated Cell Sorting (BD FACSAriaTM III, BD Biosciences, San Jose, CA, USA) with the ErbB3 membrane marker (PE anti-human erbB3/HER-3 antibody, Biolegend, #324706). The ErbB3-positive fraction was retained and further amplified in SkGM2 media. When cells were 80 to 90 % confluent, they were detached with Trypsin-EDTA for 7 min at 37°C, spun down 5 min at 300 g and resuspended in Cryostor freezing medium for long term cryopreservation in liquid nitrogen.

### Generation of engineered muscle tissues

FACS-purified ErbB3+ myogenic progenitor cells were detached from the culture plate with Trypsin-EDTA, cells were harnessed in DMEM-F12/FBS to inactivate Trypsin and spun down 5 min at 300 g and then resuspended in Matrigel (1:5 dilution in F10, Ham’s F10 nutrient mix, Thermo Fisher, #31550023), fibrinogen (1:10 dilution, fibrinogen bovine, Sigma-Aldrich, #F8630-1G) and thrombin (1:5 dilution, thrombin from bovine plasma, Sigma-Aldrich, #T4648-1KU). Engineered muscle tissues were then generated with our custom-made MyoStretch device. MyoStretch is a 3D-printed culture system consisting in two mobile carts sliding relative to a central module. Each module fits in a well of a 6-well plate and carries four rigid posts, each acting as an EMT anchor. Each mobile cart includes two flexible posts made of polydimethylsiloxane (PDMS). To ensure reliable force measurements, the deflection of each post is calibrated *in situ* by measuring the horizontal displacement induced by a series of known weights, using a smartphone camera, iV-Cam (e2eSoft) and Micro-Manager (^44^). Fully assembled and calibrated MyoStretch devices are then sterilized by incubation in 70% ethanol. For EMT production, PDMS molds were used to maintain the hydrogel containing myogenic progenitors between one rigid and one flexible posts. Gel formation was performed at 37 °C for 80 minutes and MyoStretch modules including up to 4 EMTs were then transferred in standard 6-well plates with SkGM2 media supplemented with 6-aminocaproic acid 2g/L (ACA, Sigma-Aldrich, #A2504). After one day, they were transferred to DMEM/F12, ITS 1X, 2% KSR, 1µM CHIRON and 5 µM SB431542 (STEMCELL Technologies, #72234) supplemented with ACA 2g/L for 7 days followed by ACA 5g/L. Uniaxial mechanical stretch was applied by moving the mobile carts away from the central module thanks to a dedicated screw on top of the device. The elongation was set at 15% of the EMT initial length thanks to a camera and the iVCam software (v.7.3.5). At the end of experiment, EMTs were directly used for contraction measurement or detached from the posts and frozen at -80°C for cryosection or RNA extraction.

### Contraction measurements

After 10 days of differentiation, EMTs were subjected to electrical stimulation (IonOptix stimulator, 20V) and contractile force was measured according to the displacement of the flexible post. The protocol includes conditioning stimulations at 0.5 Hz to trigger twitches, followed by a 100 Hz stimulation train during 700 ms to elicit a tetanic force response. EMTs were first incubated in Normal Physiological Solution (NPS, 140 mM NaCl, 5 mM KCl, 1 mM MgCl2 (all from VWR International), 10mM HEPES, 10 mM glucose, and 1.0 mM CaCl2 (all from Sigma-Aldrich) at pH 7.35, 37°C) and then in NPS without calcium to analyze their dependence on extracellular calcium. Acquisition and analysis were performed with the IonWizard software. A specific ImageJ-FiJi plugin was created to analyze contraction traces, and in particular to extract peak amplitude values after at 100 Hz. Contractile forces were then calculated from peak amplitude values, considering the stiffness features determined for each post prior to EMT formation (see Generation of engineered muscle tissues).

### Immunostaining and fluorescence microscopy

EMTs were fixed in 4% formaldehyde (Formaldehyde 16%, Thermo Fisher, #28908) overnight at 4°C. The following day, they were rapidly washed with PBS and incubated successively in sucrose (Sucrose, Sigma-Aldrich, #S0389) 7.5% during 2 hours at room temperature (RT), sucrose 20% 2 hours at RT, and sucrose 30% overnight at 4°C. Next, EMTs were detached from the device, rapidly washed with one drop of PBS on a glass slide, and embedded between 2 layers of Tissue freezing medium (TFM, Microm Microtech France, #TFM-5) in plastic freezer inclusion molds (Leica Biosystems, #3803025). These molds were then immersed in isopentane (2-methylbutane, Sigma-Aldrich, #M32631) cooled in liquid nitrogen. EMT cryosections were realized at the MicroPICell core facility (SFR Bonamy, BioCore, INSERM UMS 0.16, CNRS UAR 3556, Nantes, France) with a dedicated cryostat (CM1950, Leica Biosystems). Longitudinal, 10µm-thick sections were obtained at 50µm intervals and then permeabilized with phosphate buffered saline (PBS 1X, Corning, #21-040-CV), BSA 2.5% (Bovine serum albumin, Sigma-Aldrich, #A3059) and Triton 0.5% (Triton 100X, Eurobio, #GAUTTR001) for one hour at RT and incubated overnight at 4°C with an anti-α-actinin antibody (Sigma-Aldrich, #A7811) diluted at 1:500 in permeabilization solution. The sections were then washed three times with PBS and incubated with AlexaFluor555 secondary antibody (Goat anti-mouse IgG cross-adsorbed secondary antibody Alexa Fluor 555, Thermo Fisher, #A-21147) diluted at 1:1000 and DAPI (DAPI, Thermo Fisher, #D1306) diluted at 1:10000 for 45 minutes in the dark at RT. After three PBS washes, slides were mounted in ProLong Gold Antifade reagent mounting medium (Thermo Fisher, #P36934). The slides were observed under an upright epifluorescence microscope (Nikon Eclipse 90i, Nikon Instruments Inc., Tokyo, Japan) and images were acquired using NIS-elements software (version 4.60). Images were analyzed and formatted for publication using ImageJ software (ImageJ 1.54p). Myotube alignment was defined as the angular deviation of cells from the elongation axis. Data is presented as mean, and the comparison between the groups was carried out by using non-parametric Mann-Whitney tests, where p-values <0.05 were considered statistically significant. Data is presented using GraphPad Prism software (v8.0.1).

### Vector design and production

The reporter rAAV vector was produced in our internal viral vector manufacturing center (ViVeM, https://umr1089.univ-nantes.fr/en/facilities-cores/cpv). The reporter expression cassette including the murine secreted embryonic alkaline phosphatase (mSEAP) gene as a reporter downstream of the Spc5.12 promoter (^24^) was cloned between AAV2 inverted terminal repeats using DH10B competent bacteria (Thermo Fisher # EC0113), subsequently used for vector production. The vector plasmid was transfected in adherent human embryonic kidney 293 cells together with pDPx plasmids containing viral sequences required for replication and encapsidation. The plasmids containing the sequences coding for the MyoAAV variants were provided by Dr. Pardis Sabeti (Broad Institute, Cambridge, USA) after signature of a dedicated MTA. The cells were then harvested 48h to 96h after transfection, or 60h to 72h for serotype 8. The rAAV particles present in the supernatant were harvested, precipitated with PolyEthyleneGlycol and Benzonase® was added to eliminate contaminating DNA. The vectors were then purified using two successive cesium chloride gradients. The cesium chloride was removed by dialysis and the vectors were frozen at <-70°C. The particles containing the transgene were titrated by qPCR targeting the ITR2 sequences.

### rAAV transduction

rAAV vectors were diluted in DPBS and the transduction volume was adjusted to 7.5 µl / EMT. The vector mix was added during the hydrogel casting process with cells, Matrigel® and fibrinogen, before adding thrombin to activate fibrin gel formation.

### Luminometry

Culture media was collected every other day after rAAV transduction and frozen at <-70°C until analysis. Detection of the alkaline phosphatase activity was carried out with a commercially available chemiluminescence assay (Phospha-Light kit, Thermo Fisher, #T1017) following the manufacturer’s instructions. The assay is based on the use of the substrate CSPD [3-(4-methoxyspirol-[1,2-dioxetane-3,2(5’-chloro)-tricyclo(3.3.1.1. decane]-4-yl) phenyl phosphate] which is dephosphorylated by alkaline phosphatase. This results in an unstable dioxetane anion, which decomposes and emits light with its maximum activity at a wavelength of 477 nm. CSPD substrate was diluted with Reaction Buffer diluent (1:20) to obtain the reaction buffer (50µL/well). Assay buffer (50µL/well) and reaction buffer prepared above were equilibrated at RT. Samples were prepared by diluting 75µL of supernatants with 75µL of dilution buffer. Successive dilutions of the positive control (purified human placental alkaline phosphatase) were also performed in dilution buffer. The samples were heated at 65°C for 30 minutes to inactivate endogenous phosphatases and then cooled to RT. Diluted sample were transferred to opaque 96-well plates (Greiner bio-one, #655079) before addition of the assay buffer and incubation for 5 minutes at RT. The reaction buffer was then added in each well and incubated for 20 minutes at RT. Microplates were placed in a luminometer (Multimode microplate reader Spark®, Tecan) and the luminescence acquisition was performed for 1000 ms/well. The results were expressed in relative light units per second. The serial dilution of the positive control was used to validate that the samples fall within the linear range of the assay. The data was processed using GraphPad Prism software, and the associated statistical analysis were also carried out by using non-parametric Mann-Whitney tests, where p-values <0.05 were considered statistically significant.

### RTqPCR

Total RNA was purified after EMT lysis in TRIzol® (QIAzol® lysis reagent, Qiagen, #79306) and chloroform-based extraction (Chloroform, Sigma-Aldrich, #372978). Briefly, 1 mL of TRIzol was added on freshly thawed EMTs to avoid RNA degradation, together with 2 TissueLyserII beads (QIAGEN). After homogenization for 30 sec at 30 Hz, 200 μL of chloroform was added and debris were centrifuged at 12,000× g at +4 ◦C. The upper aqueous phase was collected and mixed with glycogen (Glycogen RNA grade, ThermoFischer Scientific, #R0551) and isopropanol (Propanol-2, VWR, #20842.298). After precipitation, pellets were washed with 75% ethanol and resuspended in RNAse-free H2O (Thermo Fisher). RNA samples were titrated on a Nanodrop instrument (NanoDrop One C, ThermoFischer Scientific), and then treated with RNAse-free DNAse I from the TURBO™ DNA-free Kit (ThermoFischer Scientific, #AM1907). DNA-free RNA samples were then reverse transcribed using the M-MLV RTase kit according to manufacturer recommendations (Thermo Fisher, #28025013). Transcripts were quantified using a Biorad thermocycler (CFX96™ Real-time system, Bio-Rad).

The primer used for mSEAP (Appl2) cDNA quantification were as follows: forward primer 5′-TAGTTCAAGAGTGGCTGGCA-3′, reverse primer 5′-GGGTCTCGGTGGATTTCATA-3′. The reverse-transcribed mRNA measurement was normalized by quantifying the expression of endogenous *UBC* (ubiquitin C) gene using the following primers: forward primer 5′-AGTAGTCCCTTCTCGGCGAT-3′ and reverse 5′-GCATTGTCAAGTGACGATCACAG-3′. The mSEAP and *UBC* qPCR reactions were performed with 12.5 ng of cDNA using SYBRgreen technology (TB Green Premix Ex Taq II, Takara) and used the following program: initial hold stage for 2 min at 50°C and initial denaturation for 10 min at 95°C, followed by 45 amplification cycles of 15 s at 95°C and 1 min at 60°C, and 1 melt stage of 95°C for 15 s, 60°C for 1 min and gradual rise in temperature from 60 to 95°C, at a rate of +0.3°C per cycle. The Ct results obtained for the transgene transcripts were normalized with UBC Ct values to obtain a Relative Quantity: RQ = 2^−ΔCt^ where ΔCt = Ct_transgene − Ct_endogenous. All the data was processed using GraphPad Prism software (v8.0.1), and the associated statistical analysis were carried out by using non-parametric Mann-Whitney tests, where p-values <0.05 were considered statistically significant.

## ACKNOWLEDGEMENTS

This work was funded by the INSERM ATIP Avenir program, Nantes Université, the University Hospital of Nantes and the Association Française contre les Myopathies (AFM) Téléthon. We acknowledge the IBISA MicroPICell facility (Nantes Université, SFR Bonamy, UMS Biocore, Biogenouest), member of the national infrastructure France-Bioimaging supported by the French National Research Agency (ANR-10-INBS-04), for their help with fluorescence image acquisition and analysis. We thank the Cytocell – Flow Cytometry and FACS core facility (SFR Bonamy, BioCore, INSERM UMS 016, CNRS UAR 3556, Nantes, France) for its technical expertise and help, member of the Scientific Interest Group (GIS) Biogenouest and the Labex IGO program supported by the French National Research Agency (ANR-11-LABX-0046-01). Finally, we thank Dr. Mohammadsharif Tabebordbar and Dr. Pardis C. Sabeti (Broad Institute of MIT and Harvard, Cambridge, USA) for allowing us to use the MyoAAV variants.

## AUTHOR CONTRIBUTIONS (CL, BR, MC, GM, CM, DLM, OA, CLG, BF, JBD)

Conceptualization: C.L.G., B.F., and J.B.D.; methodology: C.L., B.R., M.C., G.M., B.F., and J.B.D.; validation: C.L., B.R., M.C.; formal analysis: C.L., and J.B.D.; investigation: C.L., C.L.G., B.F., and J.B.D.; resources: C.M., D.L.M., O.A., B.F., and J.B.D.; data curation: C.L., G.M., B.F., and J.B.D., writing – original draft: C.L., C.L.G., B.F., and J.B.D.; writing – review and editing: all co-authors; visualization: C.L., B.F., and J.B.D., project administration: J.B.D.; funding acquisition: D.L.M., O.A., and J.B.D.

## DATA AVAILABILITY

The data supporting the results of this study can be obtained from the corresponding author upon reasonable request.

## SUPPLEMENTARY MATERIALS

**Figure S1:**
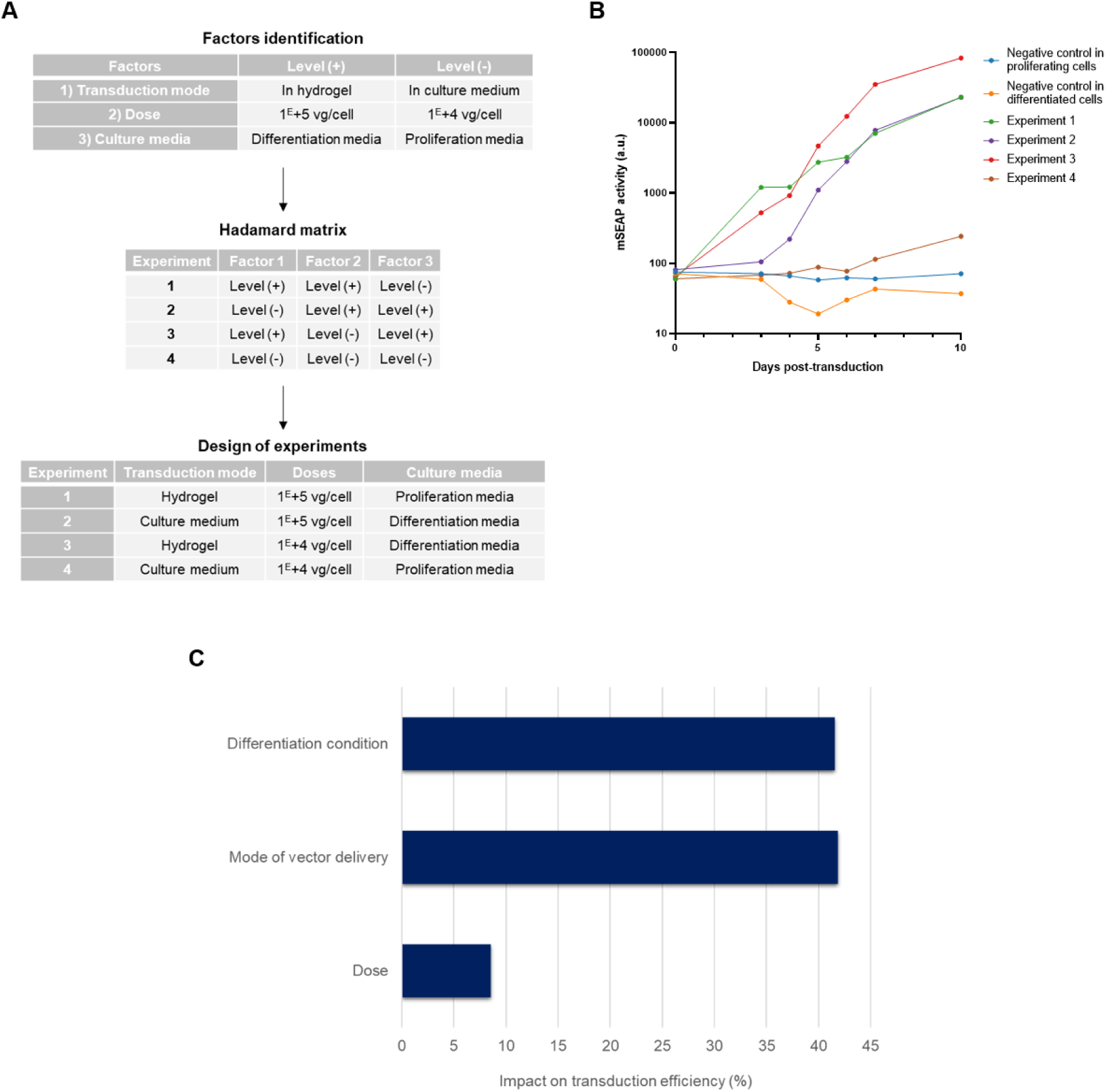
Design of experiments (DOE) approach to determine the optimal transduction conditions of iPSC-derived EMTs with reporter rAAV6-Spc5.12-mSEAP. (A) Tables summarizing the parameter combinations tested with identification of factors at 2 levels, Hadamard matrix and design of experiments. (B) Kinetics of mSEAP activity during 10 days after transduction. The luminescence values are represented in arbitrary units (a.u.) with logarithmic scale. (C) Pareto graph to determine the impact of each factor on transduction efficiency (*i.e*. transgene product expression).

**Figure S2:**
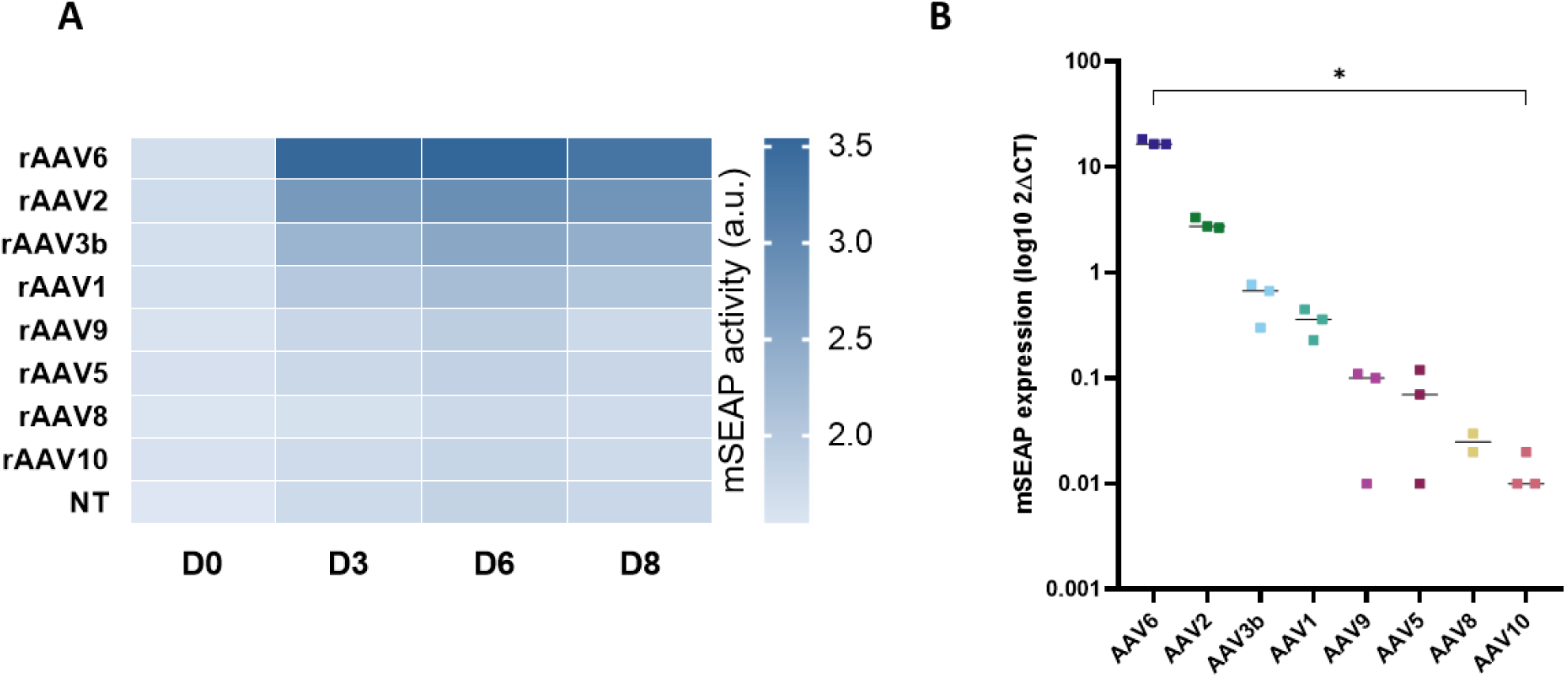
Capsid comparison in iPSC-derived myogenic progenitors in 2D differentiation. (A) Kinetics of mSEAP activity after transduction of myogenic progenitors in 2D with different serotypes of rAAV-Spc5.12-mSEAP. A darker blue box represents a higher luminescence value and therefore a higher transgene activity. The luminescence values are represented in arbitrary units (a.u.) with logarithmic scale. (B) Relative quantity of mSEAP after RNA extraction from iPSC-derived myogenic progenitors in 2D at 8 days after transduction with different rAAV serotypes. Each point represents an independent replicate (n=2 or 3 per group). Bars indicate median values. Statistics: Kruskal-Wallis test (*p-value<0.05).

**Figure S3:**
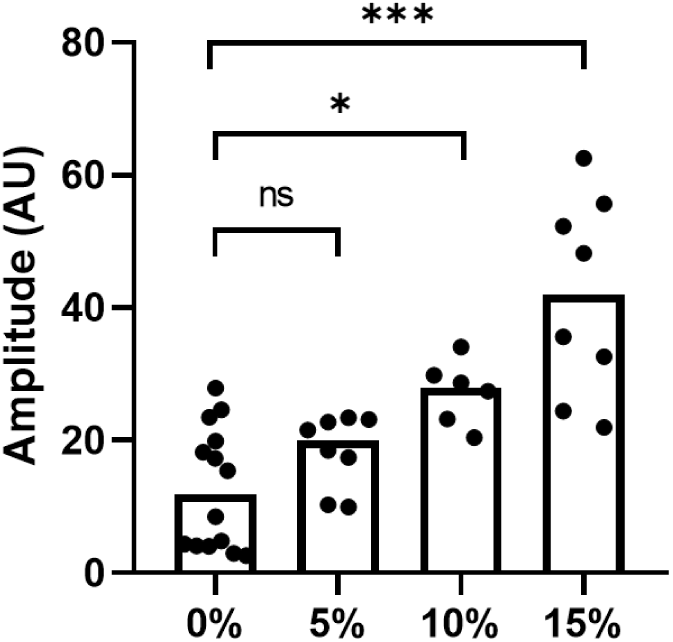
Comparison of contraction amplitude after electrical stimulation (20V, 100Hz) of EMTs stretched at +5, +10 or +15% of their initial length with unstretched EMTs. Bars indicate median. Datas were analysed by Kruskal-Wallis test (*ns* : non-significant, \**p*-value<0.05, *** *p*-value<0001).

## REFERENCES

1. Le Guiner, C., Servais, L., Montus, M., Larcher, T., Fraysse, B., Moullec, S., Allais, M., François, V., Dutilleul, M., Malerba, A., et al. (2017). Long-term microdystrophin gene therapy is effective in a canine model of Duchenne muscular dystrophy. Nat Commun 8, 16105. 10.1038/ncomms16105.

2. Le Guiner, C., Xiao, X., Larcher, T., Lafoux, A., Huchet, C., Toumaniantz, G., Adjali, O., Anegon, I., Remy, S., Grieger, J., et al. (2023). Evaluation of an AAV9-mini-dystrophin gene therapy candidate in a rat model of Duchenne muscular dystrophy. Mol Ther Methods Clin Dev 30, 30–47. 10.1016/j.omtm.2023.05.017.

3. Mack, D.L., Poulard, K., Goddard, M.A., Latournerie, V., Snyder, J.M., Grange, R.W., Elverman, M.R., Denard, J., Veron, P., Buscara, L., et al. (2017). Systemic AAV8-Mediated Gene Therapy Drives Whole-Body Correction of Myotubular Myopathy in Dogs. Mol Ther 25, 839–854. 10.1016/j.ymthe.2017.02.004.

4. Bengtsson, N.E., Tasfaout, H., and Chamberlain, J.S. (2025). The road toward AAV-mediated gene therapy of Duchenne muscular dystrophy. Mol Ther 33, 2035–2051. 10.1016/j.ymthe.2025.03.065.

5. Elangkovan, N., and Dickson, G. (2021). Gene Therapy for Duchenne Muscular Dystrophy. J Neuromuscul Dis 8, S303–S316. 10.3233/JND-210678.

6. Byrne, B.J., Butterfield, R.J., Shieh, P.B., Smith, E.C., Licht, C., Binks, M., Casinghino, S., Delnomdedieu, M., Ravindra, K.C., McDonnell, T., et al. (2025). Complement activation in a phase Ib study of fordadistrogene movaparvovec for Duchenne muscular dystrophy. Molecular Therapy 33, 4226–4238. 10.1016/j.ymthe.2025.06.032.

7. Duan, D. (2023). Lethal immunotoxicity in high-dose systemic AAV therapy. Molecular Therapy 31, 3123–3126. 10.1016/j.ymthe.2023.10.015.

8. Bönnemann, C.G., Belluscio, B.A., Braun, S., Morris, C., Singh, T., and Muntoni, F. (2023). Dystrophin Immunity after Gene Therapy for Duchenne’s Muscular Dystrophy. N Engl J Med 388, 2294–2296. 10.1056/NEJMc2212912.

9. Nathwani, A.C., Tuddenham, E.G.D., Rangarajan, S., Rosales, C., McIntosh, J., Linch, D.C., Chowdary, P., Riddell, A., Pie, A.J., Harrington, C., et al. (2011). Adenovirus-associated virus vector-mediated gene transfer in hemophilia B. N Engl J Med 365, 2357– 2365. 10.1056/NEJMoa1108046.

10. Cerneckis, J., Cai, H., and Shi, Y. (2024). Induced pluripotent stem cells (iPSCs): molecular mechanisms of induction and applications. Sig Transduct Target Ther 9, 112. 10.1038/s41392-024-01809-0.

11. Choi, I.Y., Lim, H., Estrellas, K., Mula, J., Cohen, T.V., Zhang, Y., Donnelly, C.J., Richard, J.-P., Kim, Y.J., Kim, H., et al. (2016). Concordant but Varied Phenotypes among Duchenne Muscular Dystrophy Patient-Specific Myoblasts Derived using a Human iPSC-Based Model. Cell Rep 15, 2301–2312. 10.1016/j.celrep.2016.05.016.

12. Young, C.S., Hicks, M.R., Ermolova, N.V., Nakano, H., Jan, M., Younesi, S., Karumbayaram, S., Kumagai-Cresse, C., Wang, D., Zack, J.A., et al. (2016). A Single CRISPR-Cas9 Deletion Strategy that Targets the Majority of DMD Patients Restores Dystrophin Function in hiPSC-Derived Muscle Cells. Cell Stem Cell 18, 533–540. 10.1016/j.stem.2016.01.021.

13. Al Tanoury, Z., Zimmerman, J.F., Rao, J., Sieiro, D., McNamara, H.M., Cherrier, T., Rodríguez-delaRosa, A., Hick-Colin, A., Bousson, F., Fugier-Schmucker, C., et al. (2021). Prednisolone rescues Duchenne muscular dystrophy phenotypes in human pluripotent stem cell-derived skeletal muscle in vitro. Proc Natl Acad Sci U S A 118, e2022960118. 10.1073/pnas.2022960118.

14. Sun, C., Choi, I.Y., Gonzalez, Y.I.R., Andersen, P., Talbot, C.C., Iyer, S.R., Lovering, R.M., Wagner, K.R., and Lee, G. (2020). Duchenne muscular dystrophy hiPSC–derived myoblast drug screen identifies compounds that ameliorate disease in *mdx* mice. JCI Insight 5. 10.1172/jci.insight.134287.

15. Hoch, L., Bourg, N., Degrugillier, F., Bruge, C., Benabides, M., Pellier, E., Tournois, J., Mahé, G., Maignan, N., Dawe, J., et al. (2022). Dual Blockade of Misfolded Alpha-Sarcoglycan Degradation by Bortezomib and Givinostat Combination. Front Pharmacol 13, 856804. 10.3389/fphar.2022.856804.

16. Smith, A.S., Luttrell, S.M., Dupont, J.-B., Gray, K., Lih, D., Fleming, J.W., Cunningham, N.J., Jepson, S., Hesson, J., Mathieu, J., et al. (2022). High-throughput, real-time monitoring of engineered skeletal muscle function using magnetic sensing. J Tissue Eng 13, 20417314221122127. 10.1177/20417314221122127.

17. Shahriyari, M., Islam, M.R., Sakib, S.M., Rinn, M., Rika, A., Krüger, D., Kaurani, L., Gisa, V., Winterhoff, M., Anandakumar, H., et al. (2022). Engineered skeletal muscle recapitulates human muscle development, regeneration and dystrophy. Journal of Cachexia, Sarcopenia and Muscle 13, 3106–3121. 10.1002/jcsm.13094.

18. Khodabukus, A., Prabhu, N.K., Roberts, T., Buldo, M., Detwiler, A., Fralish, Z.D., Kondash, M.E., Truskey, G.A., Koves, T.R., and Bursac, N. (2024). Bioengineered Model of Human LGMD2B Skeletal Muscle Reveals Roles of Intracellular Calcium Overload in Contractile and Metabolic Dysfunction in Dysferlinopathy. Adv Sci (Weinh) 11, 2400188. 10.1002/advs.202400188.

19. Maffioletti, S.M., Sarcar, S., Henderson, A.B.H., Mannhardt, I., Pinton, L., Moyle, L.A., Steele-Stallard, H., Cappellari, O., Wells, K.E., Ferrari, G., et al. (2018). Three-Dimensional Human iPSC-Derived Artificial Skeletal Muscles Model Muscular Dystrophies and Enable Multilineage Tissue Engineering. Cell Rep 23, 899–908. 10.1016/j.celrep.2018.03.091.

20. Pinton, L., Khedr, M., Lionello, V.M., Sarcar, S., Maffioletti, S.M., Dastidar, S., Negroni, E., Choi, S., Khokhar, N., Bigot, A., et al. (2023). 3D human induced pluripotent stem cell–derived bioengineered skeletal muscles for tissue, disease and therapy modeling. Nat Protoc 18, 1337–1376. 10.1038/s41596-022-00790-8.

21. Candiani, G., Riboldi, S.A., Sadr, N., Lorenzoni, S., Neuenschwander, P., Montevecchi, F.M., and Mantero, S. (2010). Cyclic mechanical stimulation favors myosin heavy chain accumulation in engineered skeletal muscle constructs. J Appl Biomater Biomech 8, 68–75.

22. Powell, C.A., Smiley, B.L., Mills, J., and Vandenburgh, H.H. (2002). Mechanical stimulation improves tissue-engineered human skeletal muscle. American Journal of Physiology-Cell Physiology 283, C1557–C1565. 10.1152/ajpcell.00595.2001.

23. Chal, J., Al Tanoury, Z., Hestin, M., Gobert, B., Aivio, S., Hick, A., Cherrier, T., Nesmith, A.P., Parker, K.K., and Pourquié, O. (2016). Generation of human muscle fibers and satellite-like cells from human pluripotent stem cells in vitro. Nat Protoc 11, 1833–1850. 10.1038/nprot.2016.110.

24. Li, X., Eastman, E.M., Schwartz, R.J., and Draghia-Akli, R. (1999). Synthetic muscle promoters: activities exceeding naturally occurring regulatory sequences. Nat Biotechnol 17, 241–245. 10.1038/6981.

25. Issa, S.S., Shaimardanova, A.A., Solovyeva, V.V., and Rizvanov, A.A. (2023). Various AAV Serotypes and Their Applications in Gene Therapy: An Overview. Cells 12, 785. 10.3390/cells12050785.

26. Gregorevic, P., Allen, J.M., Minami, E., Blankinship, M.J., Haraguchi, M., Meuse, L., Finn, E., Adams, M.E., Froehner, S.C., Murry, C.E., et al. (2006). rAAV6-microdystrophin preserves muscle function and extends lifespan in severely dystrophic mice. Nat Med 12, 787–789. 10.1038/nm1439.

27. Chamberlain, J.R., and Chamberlain, J.S. (2017). Progress toward Gene Therapy for Duchenne Muscular Dystrophy. Molecular Therapy 25, 1125–1131. 10.1016/j.ymthe.2017.02.019.

28. Flucher, B.E., Takekura, H., and Franzini-Armstrong, C. (1993). Development of the excitation-contraction coupling apparatus in skeletal muscle: association of sarcoplasmic reticulum and transverse tubules with myofibrils. Dev Biol 160, 135–147. 10.1006/dbio.1993.1292.

29. Weinmann, J., Weis, S., Sippel, J., Tulalamba, W., Remes, A., El Andari, J., Herrmann, A.-K., Pham, Q.H., Borowski, C., Hille, S., et al. (2020). Identification of a myotropic AAV by massively parallel in vivo evaluation of barcoded capsid variants. Nat Commun 11, 5432. 10.1038/s41467-020-19230-w.

30. Tabebordbar, M., Lagerborg, K.A., Stanton, A., King, E.M., Ye, S., Tellez, L., Krunnfusz, A., Tavakoli, S., Widrick, J.J., Messemer, K.A., et al. (2021). Directed evolution of a family of AAV capsid variants enabling potent muscle-directed gene delivery across species. Cell 184, 4919–4938.e22. 10.1016/j.cell.2021.08.028.

31. Deconinck, A.E., Rafael, J.A., Skinner, J.A., Brown, S.C., Potter, A.C., Metzinger, L., Watt, D.J., Dickson, J.G., Tinsley, J.M., and Davies, K.E. (1997). Utrophin-dystrophin-deficient mice as a model for Duchenne muscular dystrophy. Cell 90, 717–727. 10.1016/s0092-8674(00)80532-2.

32. Kipling, D., and Cooke, H.J. (1990). Hypervariable ultra-long telomeres in mice. Nature 347, 400–402. 10.1038/347400a0.

33. Chandrasekharan, K., Yoon, J.H., Xu, Y., deVries, S., Camboni, M., Janssen, P.M.L., Varki, A., and Martin, P.T. (2010). A human-specific deletion in mouse Cmah increases disease severity in the mdx model of Duchenne muscular dystrophy. Sci Transl Med 2, 42ra54. 10.1126/scitranslmed.3000692.

34. Kornegay, J.N. (2017). The golden retriever model of Duchenne muscular dystrophy. Skelet Muscle 7, 9. 10.1186/s13395-017-0124-z.

35. Kim, J., Magli, A., Chan, S.S.K., Oliveira, V.K.P., Wu, J., Darabi, R., Kyba, M., and Perlingeiro, R.C.R. (2017). Expansion and Purification Are Critical for the Therapeutic Application of Pluripotent Stem Cell-Derived Myogenic Progenitors. Stem Cell Reports 9, 12–22. 10.1016/j.stemcr.2017.04.022.

36. Hicks, M.R., Hiserodt, J., Paras, K., Fujiwara, W., Eskin, A., Jan, M., Xi, H., Young, C.S., Evseenko, D., Nelson, S.F., et al. (2018). ERBB3 and NGFR mark a distinct skeletal muscle progenitor cell in human development and hPSCs. Nat. Cell Biol. 20, 46–57. 10.1038/s41556-017-0010-2.

37. Lemke, S.B., and Schnorrer, F. (2017). Mechanical forces during muscle development. Mechanisms of Development 144, 92–101. 10.1016/j.mod.2016.11.003.

38. Vandenburgh, H.H., Hatfaludy, S., Karlisch, P., and Shansky, J. (1989). Skeletal muscle growth is stimulated by intermittent stretch-relaxation in tissue culture. Am J Physiol 256, C674–682. 10.1152/ajpcell.1989.256.3.C674.

39. Carson, J.A., and Booth, F.W. (1998). Effect of serum and mechanical stretch on skeletal α-actin gene regulation in cultured primary muscle cells. American Journal of Physiology-Cell Physiology 275, C1438–C1448. 10.1152/ajpcell.1998.275.6.C1438.

40. Ren, D., Song, J., Liu, R., Zeng, X., Yan, X., Zhang, Q., and Yuan, X. (2021). Molecular and Biomechanical Adaptations to Mechanical Stretch in Cultured Myotubes. Front. Physiol. 12. 10.3389/fphys.2021.689492.

41. Greig, J.A., Martins, K.M., Breton, C., Lamontagne, R.J., Zhu, Y., He, Z., White, J., Zhu, J.-X., Chichester, J.A., Zheng, Q., et al. (2024). Integrated vector genomes may contribute to long-term expression in primate liver after AAV administration. Nat Biotechnol 42, 1232–1242. 10.1038/s41587-023-01974-7.

42. El Andari, J., Renaud-Gabardos, E., Tulalamba, W., Weinmann, J., Mangin, L., Pham, Q.H., Hille, S., Bennett, A., Attebi, E., Bourges, E., et al. (2022). Semirational bioengineering of AAV vectors with increased potency and specificity for systemic gene therapy of muscle disorders. Sci Adv 8, eabn4704. 10.1126/sciadv.abn4704.

43. Guan, X., Mack, D.L., Moreno, C.M., Strande, J.L., Mathieu, J., Shi, Y., Markert, C.D., Wang, Z., Liu, G., Lawlor, M.W., et al. (2014). Dystrophin-deficient cardiomyocytes derived from human urine: new biologic reagents for drug discovery. Stem Cell Res 12, 467–480. 10.1016/j.scr.2013.12.004.

44. Edelstein, A.D., Tsuchida, M.A., Amodaj, N., Pinkard, H., Vale, R.D., and Stuurman, N. (2014). Advanced methods of microscope control using μManager software. Journal of Biological Methods 1, 1. 10.14440/jbm.2014.36.

