## Supplemental Material for "Engineered muscle tissues with enhanced maturation enable the identification of clinically relevant rAAV capsids"

**A**

**Factors identification**

| Factors | Level (+) | Level (-) |
|---|---|---|
| 1) Transduction mode | In hydrogel | In culture medium |
| 2) Dose | 1E+5 vg/cell | 1E+4 vg/cell |
| 3) Culture media | Differentiation media | Proliferation media |

↓

**Hadamard matrix**

| Experiment | Factor 1 | Factor 2 | Factor 3 |
|---|---|---|---|
| 1 | Level (+) | Level (+) | Level (-) |
| 2 | Level (-) | Level (+) | Level (+) |
| 3 | Level (+) | Level (-) | Level (+) |
| 4 | Level (-) | Level (-) | Level (-) |

↓

**Design of experiments**

| Experiment | Transduction mode | Doses | Culture media |
|---|---|---|---|
| 1 | Hydrogel | 1E+5 vg/cell | Proliferation media |
| 2 | Culture medium | 1E+5 vg/cell | Differentiation media |
| 3 | Hydrogel | 1E+4 vg/cell | Differentiation media |
| 4 | Culture medium | 1E+4 vg/cell | Proliferation media |

**B**

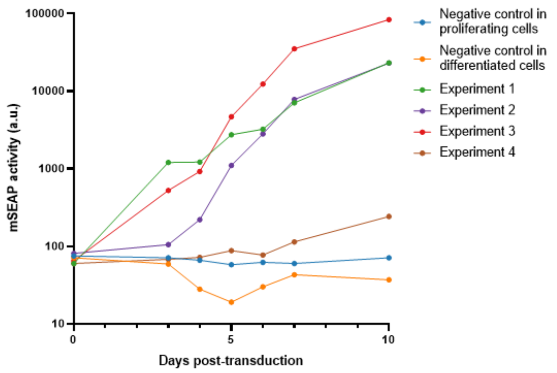


**C**

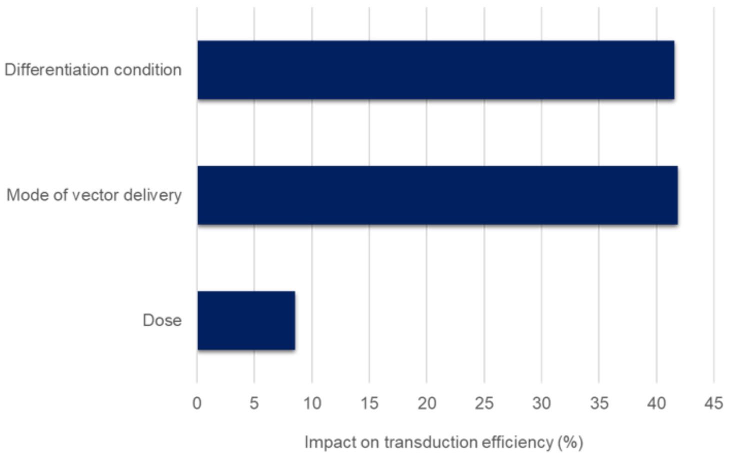


**Figure S1:** Design of experiments (DOE) approach to determine the optimal transduction conditions of iPSC-derived EMTs with reporter rAAV6-Spc5.12-mSEAP. (A) Tables summarizing the parameter combinations tested with identification of factors at 2 levels, Hadamard matrix and design of experiments. (B) Kinetics of mSEAP activity during 10 days after transduction. The luminescence values are represented in arbitrary units (a.u.) with logarithmic scale. (C) Pareto graph to determine the impact of each factor on transduction efficiency (*i.e*. transgene product expression).

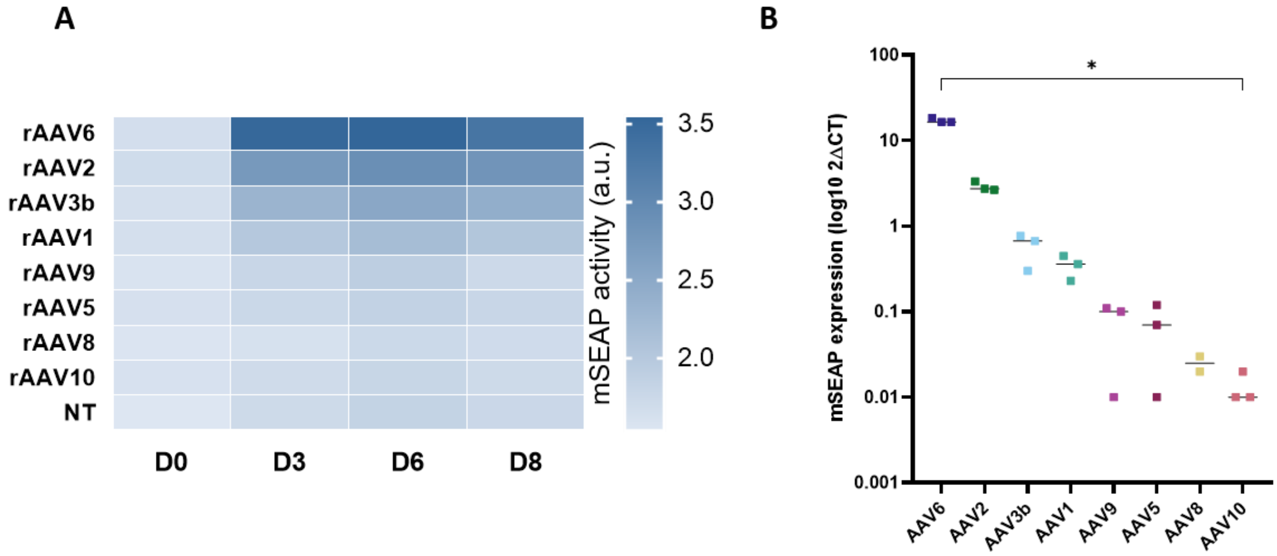


**Figure S2:** Capsid comparison in iPSC-derived myogenic progenitors in 2D differentiation. (A) Kinetics of mSEAP activity after transduction of myogenic progenitors in 2D with different serotypes of rAAV-Spc5.12-mSEAP. A darker blue box represents a higher luminescence value and therefore a higher transgene activity. The luminescence values are represented in arbitrary units (a.u.) with logarithmic scale. (B) Relative quantity of mSEAP after RNA extraction from iPSC-derived myogenic progenitors in 2D at 8 days after transduction with different rAAV serotypes. Each point represents an independent replicate (n=2 or 3 per group). Bars indicate median values. Statistics: Kruskal-Wallis test (*p-value<0.05).

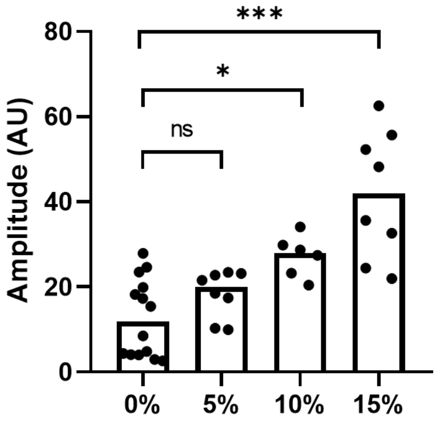


**Figure S3:** Comparison of contraction amplitude after electrical stimulation (20V, 100Hz) of EMTs stretched at +5, +10 or +15% of their initial length with unstretched EMTs. Bars indicate median. Datas were analysed by Kruskal-Wallis test (*ns* : non-significant, **p*-value<0.05, *** *p*-value<0001).
